# Mutation of Ebola virus VP35 Ser129 uncouples interferon antagonist and replication functions

**DOI:** 10.1101/726935

**Authors:** MJ Morwitzer, A Corona, L Zinzula, E Fanunza, C Nigri, S Distinto, C Vornholt, V Kumar, E Tramontano, SP Reid

## Abstract

Ebolaviruses are non-segmented, negative-sense RNA viruses (NNSVs) within the order *Mononegavirales* that possess the multifunctional virion protein 35 (VP35), a major determinant of virulence and pathogenesis that is indispensable for viral replication and host innate immune evasion. VP35 is functionally equivalent to the phosphoprotein (P) of other mononegaviruses such as rhabdoviruses and paramyxoviruses. Phosphorylation of the P protein is universally regarded as functionally important however, a regulatory role(s) of phosphorylation on VP35 function remains unexplored. Here, we identified a highly conserved Ser129 residue near the homo-oligomerization coiled coil motif, which is essential for VP35 functions. Affinity-purification MS followed by post-translational modification (PTM) analysis predicted phosphorylation of Ser129. Co-immunoprecipitation, cross-linking, and biochemical characterization studies revealed a moderately decreased capacity of VP35-S129A to oligomerize. Functional analysis showed that Ser-to-Ala substitution of Ebola virus (EBOV) VP35 did not affect IFN inhibitory activity but nearly abolished EBOV minigenome activity. Further coimmunoprecipitation studies demonstrated a lost interaction between VP35-S129A and the amino terminus of the viral polymerase but not between viral nucleoprotein (NP) or VP35-WT. Taken together, our findings provide evidence that phosphorylation modulates VP35 function, supporting VP35 as a NNSV P protein and providing a potentially valuable therapeutic target.

**Importance:** Ebola virus (EBOV) can cause severe disease in humans. The 2013-2016 West African epidemic and the two recent outbreaks in the Democratic Republic of the Congo underscore the urgent need for effective countermeasures, which remain lacking. A better understanding of EBOV biology and the modulation of multifunctional viral proteins is desperately needed to develop improved therapeutics. We provide evidence here that function of virion protein 35 (VP35) is modulated by phosphorylation of Ser129, a conserved residue among other ebolavirus species. These findings shed light on EBOV biology and present a potential target for broad acting anti-ebolavirus therapeutics.

## Introduction

Ebolaviruses are zoonotic pathogens associated with severe hemorrhagic disease in humans (Messaoudi et al., 2015, Nelson et al., 2017). The genus *Ebolavirus*, which is of the family *Filoviridae* and order *Mononegavirales*, includes six species of filamentous, enveloped, non-segmented negative-sense RNA viruses (NNSVs): Ebola (EBOV), Sudan (SUDV), Reston (RESTV), Tai Forest (TAFV), Bundibugyo (BDBV), and Bombali (BOMV) viruses (Feldmann et al., 2013, Kuhn, 2017, Goldstein et al., 2018). Among these, EBOV has caused the most substantial outbreaks, along with SUDV and BDBV (Messaoudi et al., 2015). Once sporadic, EBOV outbreaks have become increasingly frequent and large, as evidenced by the unprecedented 2013-2016 West African epidemic and the 2017 and currently ongoing outbreaks in the Democratic Republic of the Congo (WHO, 2016, CDC, 2018). Even with several promising therapeutic candidates under investigation, there remains no licensed therapeutics available to prevent or treat infection (Keshwara et al., 2017, Cross et al., 2018, Fanunza et al., 2018a). Thus, continued efforts to identify and develop therapeutics, particularly those with pan-filovirus efficacy, should be a priority.

The ebolavirus genome is approximately 19 kb and encodes nine proteins, among which the multifunctional virion protein 35 (VP35) is a major determinant of virulence and pathogenesis (Leung et al., 2010b, Feldmann et al., 2013, Messaoudi et al., 2015). VP35 is a potent immune antagonist, inhibiting interferon-α/β (IFN-α/β) production, activation of the IFN-inducible protein kinase R (PKR) antiviral protein, and RNA interference (Basler et al., 2000, Basler et al., 2003, Feng et al., 2007, Schumann et al., 2009, Haasnoot et al., 2007, Fabozzi et al., 2011). VP35 inhibits retinoic acid-inducible gene I (RIG-I)-like receptor signaling and IFN-α/β production by several mechanisms, including sequestering immunostimulatory double-stranded RNA (dsRNA) intermediates, inhibiting PACT-induced RIG-I ATPase activity, preventing kinases TANK-binding kinase 1 (TBK1), and/or IB kinase epsilon (IKKɛ) from activating interferon regulatory factors 3 and 7 (IRF3/IRF7), inactivating IRF7 by SUMOylation, and inhibiting TRIM6-mediated type I IFN production (Cardenas et al., 2006, Bale et al., 2013, Dilley et al., 2017, Luthra et al., 2013, Prins et al., 2009, Hartman et al., 2008, Chang et al., 2009, Bharaj et al., 2017). In addition to immune suppression, VP35 also functions as an essential cofactor of the viral RNA-dependent RNA polymerase (named L for large polymerase protein). The functional viral polymerase complex is comprised of nucleoprotein (NP) and VP30 in addition to L and VP35 (Muhlberger et al., 1998, Muhlberger et al., 1999, Prins et al., 2010). VP35 is thought to act as a bridge between the catalytic subunit of L and the NP-associated viral RNA, thus VP35-L and VP35-NP interactions are likely both essential for viral RNA synthesis (Becker et al., 1998, DiCarlo et al., 2007, Trunschke et al., 2013, Kirchdoerfer et al., 2015, Leung et al., 2015). Moreover, VP35 serves as a structural protein that is necessary, along with NP and VP24, for nucleocapsid formation (Huang et al., 2002, Shi et al., 2008, Takamatsu et al., 2018). Furthermore, a recent study identified novel NTPase and helicase-like activities of EBOV VP35, which suggests VP35 may also participate in viral RNA remodeling activity (Shu et al., 2019). The N-terminus of VP35 contains a NP-chaperoning domain as well as a homo-oligomerization domain, and the C-terminus contains a dsRNA/IFN inhibitory domain (RBD/IID) (Kirchdoerfer et al., 2015, Leung et al., 2015, Reid et al., 2005, Bale et al., 2012, Bale et al., 2013, Kimberlin et al., 2010, Leung et al., 2009a, Leung et al., 2009b, Leung et al., 2010c, Leung et al., 2010a, Leung et al., 2010d, Ramanan et al., 2012).Homo-oligomeric organization of VP35 occurs via a predicted coiled-coil motif in the homo-oligomerization domain, and is known to be required for replication and full IFN antagonism (Reid et al., 2005, Moller et al., 2005). All ebolavirus VP35s form tetramers, whereas only the oligomerization domain of EBOV VP35 (eVP35) is able to form trimers (Zinzula et al., 2019b, Chanthamontri et al., 2019).

Filovirus VP35 is the functional equivalent to the phosphoprotein (P) of other members of the order *Mononegavirales*, which also includes families such as *Rhabdoviridae*, *Paramyxoviridae*, and *Bornaviridae*. As its name suggests, P protein is the most highly phosphorylated viral protein in any infected cell or virion of mononegaviruses. Phosphorylation of P protein is universally regarded as functionally important, with increasing experimental support. Thus far, phosphorylation has been shown to modulate P protein function of vesicular stomatitis virus (VSV) (Chattopadhyay and Banerjee, 1987, Barik and Banerjee, 1991, Barik and Banerjee, 1992a, Barik and Banerjee, 1992b, Gao and Lenard, 1995, Gao et al., 1996, Pattnaik et al., 1997, Das and Pattnaik, 2004), respiratory syncytial virus (RSV) (Barik et al., 1995, Asenjo and Villanueva, 2000, Villanueva et al., 2000, Asenjo et al., 2006, Asenjo et al., 2008, Asenjo and Villanueva, 2016), Chandipura virus (CHPV) (Chattopadhyay et al., 1997, Raha et al., 1999, Raha et al., 2000), Borna disease virus (BDV) (Schmid et al., 2007), parainfluenza virus 5 (PIV5) (Timani et al., 2008), Rabies virus (RABV) (Moseley et al., 2007), Rinderpest (RPV) (Saikia et al., 2008), measles virus (MV)(Sugai et al., 2012), mumps virus (MuV) (Pickar et al., 2014), and Newcastle disease virus (NDV) (Qiu et al., 2016). However, it is unknown whether the function of filovirus P protein is regulated by phosphorylation. Of note, although all P proteins are thought to perform similar transcriptional and replication functions, there is surprisingly little sequence homology, even between the Indiana and New Jersey serotypes of VSV. EBOV VP35 is known to be moderately phosphorylated (Elliott et al., 1985, Prins et al., 2009). Accordingly, it might be expected that phosphorylation of VP35 is functionally important.

In this study, we aimed to further elucidate modulation of VP35 function by evaluating it as a NNSV P protein. We used affinity purification-mass spectrometry (AP-MS) to identify putative post-translational modifications (PTMs) of VP35. Phosphorylation was predicted for a highly conserved Ser129 residue proximal to the homo-oligomerization domain. Oligomerization studies indicated that an alanine mutant VP35-S129A had moderately reduced ability to homo-oligomerize. Further functional studies revealed that while the mutant retained IFN antagonist activity, minigenome activity was nearly abolished. VP35-NP and VP35–L interactions were interrogated by co-immunoprecipitation (co-IP) and immunofluorescence (IF) experiments. While VP35-NP interaction remained intact, VP35-L was abrogated, indicating that phosphorylation of VP35 at S129 is crucial to the ability of VP35 to interact with L. These results support VP35 as a NNSV P protein and further elucidate the regulation of VP35 functions, potentially offering a novel pan-filovirus therapeutic target.

## Methods

### Antibodies

Mouse anti-HA (H3663), rabbit anti-HA (H6908), mouse anti-FLAG (F3165), rabbit anti-FLAG (F7425), rabbit anti-ISG56 (SAB1410690), and mouse anti-Calnexin (C7617) antisera were purchased from Sigma-Aldrich (ST. Louis, MO, USA). Rabbit anti-VP35 (GTX134032) was purchased from GeneTex (Irvine, CA, USA). Mouse anti-His (TA150088) was purchased from OriGene (Rockville, MD, USA). Rabbit anti-L was purchased from IBT BioServices (Rockville, MD, USA).Goat anti-rabbit HRP (65-6120), goat anti-mouse HRP (32430), goat anti-mouse Alexa Fluor 488 (A32723), goat anti-rabbit Alexa Fluor 568 (A11011), and Hoechst 33342 (H3570) were purchased from ThermoFisher Scientific (Grand Island, NY, USA).

### Cell lines and viral RNA production

HeLa, HEK293T, and A549 cells were grown in Dulbecco’s modified Eagle’s medium (Gibco) supplemented with 10% fetal bovine serum (Gibco) and 1% penicillin/streptomycin (Sigma). Cells were maintained at 37°C in a humidified 5% CO_2_ incubator. Viral RNA (vRNA) was produced in A549 cells, infected with IAV/Puerto-Rico/8/34 (IAV PR8) strain at a multiplicity of infection of 5. Five hours after infection, total RNA was isolated using the RNeasy® Kit from Qiagen (Hilden, Germany). TRIzol™ Reagent was purchased from Thermo Fisher Scientific.

### Plasmids and Reagents

Plasmid pGL-IFN-β-luc was kindly provided by Professor Stephan Ludwig, Institute of Molecular Virology, University of Münster, Germany). pRL-TK was purchased form Promega (Promega Italia S.r.l. Milan, Italy). pcDNA3-EBOV VP35, carrying EBOV VP35 gene form Zaire species (1976 Yambuku-Mayinga strain) was constructed as previously reported (Cannas et al., 2015). The VP35-S129A substitution was introduced in the pcDNA3-EBOV VP35 plasmid using the QuikChange mutagenesis kit by following the manufacturer’s instructions (Agilent Technologies Inc., Santa Clara, CA). Primers’ sequences: VP35-S129A Forward:5’-GATATGGCAAAAACAATCGCCTCATTGAACAGGGTTTG-3’; VP35-S129A reverse: 5’-CAAACCCTGTTCAATGAGGCGATTGTTTTTGCCATATC-3’. pCAGGS-HA-EBOV-NP was purchased from BEI (NR-49343). pCAGGS-NP-V5, pCAGGS-V5-VP30, and pcDNA3 vectors were generated at the USAMRIID. pCAGGS-FLAG-VP35 and pCAGGS-L_1-505 were a kind gift from Dr. Christopher F. Basler (Georgia State University). pCAGGS_L_EBOV and pCAGGS_3E5E_luciferase were gifts from Dr. Elke Mühlberger (Addgene plasmid # 103052; # 103055) (Nelson et al., 2017).

jetPRIME transfection reagent was purchased from Polyplus-transfection (S.A., Illkirch, France).T-Pro P-Fect Transfection Reagent was purchased from T-Pro Biotechnology (New Taipei County, Taiwan). Lipofectamine 2000 and Lipofectamine 3000 were purchased from Thermo Fisher Scientific. D-luciferin and Coelenterazine were purchased form Gold Biotechnology (U.S. Registration No 3,257,927). Primers were purchased form (Metabion, Planegg, Germany). The Dual-Glo Luciferase Assay System (E2920) was purchased from Promega (Madison, WI, USA) and used per manufacturer’s instructions.

### Viral sequences

The following filovirus sequences were used for sequence comparison and/or cloning where indicated: EBOV (*Zaire ebolavirus* isolate Ebola virus/H.sapiens-tc/COD/1976/Yambuku-Mayinga, GenBank: NC_002549.1), SUDV (*Sudan ebolavirus* isolate Sudan virus/H.sapiens-tc/UGA/2000/Gulu-808892, GenBank: NC_006432.1), RESTV (*Reston ebolavirus* isolate Reston virus/M.fascicularis-tc/USA/1989/Philippines89-Pennsylvania, GenBank: NC_004161.1), TAFV (*Tai Forest ebolavirus* isolate Tai Forest virus/H.sapiens-tc/CIV/1994/Pauleoula-CI, GenBank: NC_014372.1), BDBV (*Bundibugyo ebolavirus* isolate Bundibugyo virus/H.sapiens-tc/UGA/2007/Butalya-811250, GenBank: NC_014373.1), BOMV (*Bombali ebolavirus* isolate Bombali ebolavirus/Mops condylurus/SLE/2016/PREDICT_SLAB000156, GenBank: NC_039345.1). Multiple sequence alignment was made by Clustal Omega (Sievers and Higgins, 2018).

### Mass spectrometry analysis

Affinity-tagged VP35 was overexpressed in HeLa cells using Lipofectamine 2000 as per manufacturer’s instruction. Affinity purified material was resolved by SDS-PAGE and stained with Coomassie (ThermoFisher Scientific). The stained protein band corresponding to protein of interest was excised and processed for in-gel digestion at the UNMC proteomics facility using the protocol of Shevchenko and colleagues (Shevchenko et al., 2006). Extracted peptides were re-suspended in 2% acetonitrile (ACN) and 0.1% formic acid (FA) and loaded onto trap column Acclaim PepMap 100 75μm × 2 cm C18 LC Columns (Thermo Scientific™) at flow rate of 4 μl/min then separated with a Thermo RSLC Ultimate 3000 (Thermo Scientific™) on a Thermo Easy-Spray PepMap RSLC C18 75μm × 50cm C-18 2 μm column (Thermo Scientific™) with a step gradient of 4–25% solvent B (0.1% FA in 80 % ACN) from 10-57 min and 25–45% solvent B for 57–62 min at 300 nL/min and 50oC with a 90 min total run time. Eluted peptides were analyzed by a Thermo Orbitrap Fusion Lumos Tribrid (Thermo Scientific™) mass spectrometer in a data dependent acquisition mode. A survey full scan MS (from m/z 350–1800) was acquired in the Orbitrap with a resolution of 120,000. The AGC target for MS1 was set as 4 × 105 and ion filling time set as 100 ms. The most intense ions with charge state 2-6 were isolated in 3 s cycle and fragmented using HCD fragmentation with 40 % normalized collision energy and detected at a mass resolution of 30,000 at 200 m/z. The AGC target for MS/MS was set as 5 × 104 and ion filling time set 60 ms dynamic exclusion was set for 30 s with a 10 ppm mass window. Protein identification was performed by searching MS/MS data against the swiss-prot human protein database downloaded on Aug 20, 2018. The search was set up for full tryptic peptides with a maximum of two missed cleavage sites. Acetylation of protein N-terminus and oxidized methionine were included as variable modifications and carbamidomethylation of cysteine was set as fixed modification. The precursor mass tolerance threshold was set 10 ppm for and maximum fragment mass error was 0.02 Da. Qualitative analysis was performed using PEAKS 8.5 software. The significance threshold of the ion score was calculated based on a false discovery rate of ≤ 1%.

### Co-immunoprecipitations (co-IPs)

HeLa cells (1 × 10^6^ cells) were transfected with the indicated plasmids using 3 μL of the transfection reagent jetPRIME (Polyplus) per 1 μg DNA per manufacturer’s instructions. The total amount of DNA for each transfection was kept constant in each experiment by complementing with empty vector. Twenty-four h post-transfection, cells were lysed in a modified RIPA buffer (50 mM Tris-HCl pH 7.4, 150 mM NaCl, 1mM EDTA, 1% NP-40, 0.25% Na-deoxycholate) containing protease and a phosphatase inhibitor cocktail (Thermo Scientific, Waltham, MA, USA). Approximately 10% WCL was reserved for IB analysis before performing co-IPs. To immunoprecipitate indicated proteins, lysates were incubated with EZview Red ANTI-FLAG M2 Affinity Gel (Sigma) or EZview Red Anti-HA Affinity Gel (Sigma) for 1 h at 4 °C. Beads were washed 4 to 5 times with TBS, re-suspended in 2X Lane Marker Reducing Sample Buffer (Thermo Scientific), and then subjected to SDS-PAGE and immunoblotting, as indicated below.

### Immunoblotting (IB)

Cell lysates were subjected to reducing SDS-PAGE and proteins were transferred onto PVDF membranes. Blots were probed with indicated primary antibodies either 1–2 h at RT or overnight at 4°C. Secondary incubations were performed for 1–2 h at RT using either goat anti-rabbit HRP or goat anti-mouse HRP. Radiance chemiluminescence substrate (Azure Biosystems; Dublin, CA, USA) was used to visualize protein on an Azure c600 imaging system.

### DSP Crosslinking

HeLa cells were transfected with VP35-WT or VP35-S129A plasmids as previously indicated. Twenty-four h post-transfection the cells were incubated in a PBS reaction buffer with or without the Dithiobis(succinimidylpropionate) DSP cross-linker (Thermo Scientific) at final concentration of 1 mM for 30 min at RT. The reaction was quenched by the addition of a stop solution (1M Tris-HCl pH 7.5) for 15 min. The cells were subsequently lysed in the modified RIPA buffer and resuspended in a non-reducing sample buffer (Thermo Scientific) and subjected to SDS-PAGE and immunoblot analysis.

### Molecular cloning, expression and purification of recombinant proteins

Constructs encoding for the recombinant EBOV (Zaire ebolavirus, Yambuku-Mayinga, GenBank: NC_002549.1) WT and S129A VP35 oligomerization domains (residues 83-175) were subcloned into pET21b vectors (Novagen) from synthetic DNA preparations (BioCat), then expressed in *E.coli* BL21 (DE3) cells and purified by affinity and size-exclusion chromatography as previously described (Zinzula et al 2019). Briefly, heat-shock transformed bacterial cells were grown in LB medium (Carl-Roth) at 37 °C and 200 rpm up to 0.8 optical density at 600 nm, then induced at 20 °C with 0.5 mM isopropyl b-D-1-thiogalactopyranoside (IPTG) (Carl-Roth) for 16 hours. Proteins were purified by affinity on a 1 mL complete HisTag Purification Column (Roche) and by size-exclusion on a Superose 12 10/300 GL (GE Healthcare) column, respectively.

### Miniaturized differential scanning fluorimetry (nanoDSF)

NanoDSF was performed by using a Prometheus NT.48 nanoDSF instrument (NanoTemper Technologies). Recombinant EBOV VP35 WT and S129A oligomerization domains were diluted at 2.5 mg/mL in 25 mM tris-HCl pH 7.4, 150 mM NaCl, 2 mM MgCl2 and loaded on 10 μL standard UV capillaries. Measurement of the instrinsic protein fluorescence at 330 and 350 nm wavelength was performed over a 20-95 °C thermal gradient run at 0.5 °C/min rate. Melting temperature (Tm) values were determined as the first derivative maxima of the intrinsic fluorescence 330/350 nm ratio functions. Data were averaged from at least three independent experiments.

### Circular dichroism (CD) spectroscopy and secondary structure content analysis

CD spectroscopy was performed on a J-715 spectropolarimeter (Jasco) flushed with N_2_ and equipped with a PDF 3505 Peltier thermostat (Jasco). Purified recombinant EBOV VP35 WT and S129A oligomerization domains were diluted at 0.1 mg/mL in 25 mM tris-HCl pH 7.4, 150 mM NaCl, 2 mM MgCl_2_ and transferred to a high-precision 1-mm pathlength quartz cuvette (Helma Analytics). Wavelength scans were recorded between 190 and 250 nm, at 4 °C, 23 °C and 37 °C, at 50 nm/min scanning speed with 1 s response time and 0.1 nm data pitch, each spectrum being the average of four accumulations. Mean molar residue ellipticity (MRE) values were calculated as MRE = [θ] = 3300 *m*ΔA/*lcn*, where *m* is the molecular mass in Daltons, A is the absorbance, *l* is the path length in centimeters, c is the protein concentration expressed in milligrams per milliliter and *n* is the number of amino acid residues. Data processing for secondary structure content determination was performed by using the CONTINLL deconvolution method and the SMP56 reference set implemented in the Spectra Manager software package.

### Size exclusion chromatography (SEC) and multi-angle light scattering (MALS)

SEC-MALS was performed on a Superdex 200 10/300 GL gel filtration column (GE Healthcare), connected to a 1100 HPLC system with variable UV absorbance detector set at 280 nm (Agilent Technologies) and coupled in line with a mini DAWN TREOS MALS detector and an Optilab rEX refractive-index detector (Wyatt Technology, 690 nm laser). Purified recombinant EBOV VP35 WT and S129A oligomerization domains (~50 to 100 μg) were loaded by autoinjection and run at 23 °C and 0.5 mL/min flow rate in 25 mM tris-HCl pH 7.4, 150 mM NaCl, 2 mM MgCl_2_. Protein absolute molecular masses were calculated with the ASTRA 6 software (Wyatt Technology) with the dn/dc value set to 0.185 mL/g. Bovine serum albumin (ThermoFisher) was used as calibration standard. Data were averaged from at least two independent experiments.

### Computational modeling

The coordinates for the oligomerization domain of VP35 from Ebola virus were taken from the RCSB Protein Data Bank (Berman et al., 2000) as reported in the entry 6GBO.(Zinzula et al., 2019a) The structure processing and optimization were carried out by means of Maestro Protein Preparation Wizard default setting (2018). Original water molecules were removed. After preparation, the structure was refined to optimize the hydrogen bond network using OPLS_2005 force field.(Kaminski et al., 2001). S129A mutation was generated starting from VP35-WT. Therefore, the mutant geometry was optimized with a full minimization protocol considering OPLS_2005 force field (Kaminski et al., 2001) and GB/SA implicit water, setting 10000 steps interactions analysis with Polak-Ribier Coniugate Gradient (PRCG) method and a convergence criterion of 0.1 kcal/(molÅ).

### Luciferase reporter gene assay for IFN promoter activation

The luciferase reporter gene assay for IFN promoter activation was adapted from (Fanunza et al., 2018b). HEK293T cells (1.5 × 10^4^ cells/well) were seeded in 96-well plates 24 h before transfection. Cells were transfected using T-Pro P-Fect Transfection Reagent, according to the manufacturer’s protocol. Plasmids pGL-IFN-β-luc (60 ng), pRL-TK (10 ng), and various dilutions (100, 10, 1.0, 0.1, 0.01 ng) of pcDNA3 vector control (VC), pcDNA3_VP35-WT, or pcDNA3_VP35-S129A were mixed with the transfection reagent in reduced serum medium Optimem (Gibco) and incubated 20 min at RT. Transfection complexes were then gently added into individual wells of the 96-well plate. Twenty-four hours after transfection, cells were stimulated with influenza A virus (IAV)-RNA, pre-mixed with the transfection reagent in reduced serum medium Optimem, and incubated for 24 h at 37 °C in 5% CO_2_. Cells were harvested with lysis buffer (50 mM Na-MES [pH 7.8], 50 mM Tris-HCl [pH 7.8], 1 mM dithiothreitol, and 0.2% Triton X-100). To lysates were added luciferase assay buffer (125 mM Na-MES [pH 7.8], 125 mM Tris-HCl [pH 7.8], 25 mM magnesium acetate, and 2.5 mg/mL ATP). Immediately after addition of 50 μl of Luciferin buffer (25 mM D-luciferin, 5 mM KH2PO4) the luminescence was measured in Victor3 luminometer (Perkin Elmer), and again after addition of 50 μ of Coelenterazine assay buffer (125 mM Na-MES [pH 7.8], 125 mM Tris-HCl [pH 7.8], 25 mM magnesium acetate, 5 mM KH2PO4 and 10 μM Coelenterazine). Relative light units (RLU) of firefly luciferase signal were normalized to *Renilla* luciferase, with percentage of IFN induction values calculated relative to unstimulated controls (indicated as 100%). Each assay was performed in triplicate. Mean ± standard deviation (SD) values and paired, two-tailed t tests were calculated using GraphPad Prism 6.01 software (GraphPad Software, Inc.; San Diego, CA, USA).

### RT-PCR assay for ISG expression

HEK293T cells (3 × 10^5^ cells/well) resuspended Dulbecco’s modified Eagle’s medium supplemented with 10% fetal bovine serum were seeded in 6-well plates, pre-treated with 500 μl of Poly-D-lysine hydrobromide 100μg/ml for 1 h. After 24 h cells were transfected with 2500 ng of plasmid (pcDNA3 vector control (VC), pcDNA3_VP35-WT, or pcDNA3_VP35-S129A) using Lipofectamine 3000 transfection reagent, as per manufacturer’s instruction. Twenty-four h after transfection, cells were stimulated with 2500 ng of IAV-RNA, pre-mixed with the transfection reagent in reduced serum medium Optimem, and incubated for 24 hours at 37 °C in 5% CO_2_. Total RNA was extracted from transfected cells with TRIzol® Reagent (Invitrogen). RNA was then reverse transcribed and amplified using Luna Universal One-Step RT- qPCR kit (New England Biolabs; Ipswich, MA, USA). Quantitative real-time PCR (RT-qPCR) experiments were performed in triplicate in a CFX-96 Real-Time system (BioRad; Hercules, CA, USA). Primers used: GAPDH Forward 5’-GAGTCAACGGATTTTGGTCGT-3’, GAPDH Reverse 5’-TTGATTTTGGAGGGATCTCG-3’; ISG15 Forward 5’-TCCTGGTGAGGAATAACAAGGG-3’; ISG15 Reverse 5’-CTCAGCCAGAACAGGTCGTC-3’; 2’5’OAS Forward 5’-AGCTTCATGGAGAGGGGCA-3’; 2’5’OAS Reverse 5’-AGGCCTGGCTGAATTACCCAT-3’. Data were analyzed with Opticon Monitor 3.1. mRNA expression levels were normalized to the level of glyceraldehyde-3-phosphate dehydrogenase (GAPDH), with percentage of gene expression over the VC (indicated as 100%). Mean ± standard deviation values of two replicates and paired, two-tailed t tests were calculated using GraphPad Prism 6.01 software.

### Immunoblot for ISG56 expression

HeLa cells (1 × 10^6^ cells) were transfected with the indicated plasmids using 2 μL of the transfection reagent Lipofectamine 2000 (Thermo Fisher Scientific) per 1 μg DNA per manufacturer’s instructions. Twenty four h post-transfection the cells were mock treated or treated with 10 μg/mL of poly I:C (InvivoGen). Twenty-four h post -treatment cells were harvested, lysed in modified RIPA buffer and subjected to SDS-PAGE and immunoblot as aforementioned with the indicated antibodies.

### Minigenome Assay

A RNA polymerase II-driven EBOV minigenome was used as previously described (Nelson et al., 2017). Briefly, HeLa cells (3.0 × 10^5^) were seeded in 12-well plates 24 h before transfection. Cells were transfected with minigenome components (125 ng pCAGGS-HA-NP, 125 ng pCAGGS-FLAG-VP35, 50 ng pCAGGS-V5-VP30, 500 ng pCAGGS-L, and 750 ng of pCAGGS-3E5E-luciferase) along with 50 ng pRL-TK (for transfection efficiency control) using jetPRIME reagent as per manufacturer’s recommendation. For the no L control, total DNA levels were kept constant by complementing transfections with empty-vector pcDNA3. Reporter activity was measured 48 h post-transfection using the Dual-Glo Luciferase Assay System and a Tecan Spark microplate luminometer (Tecan Trading AG, Switzerland). Whole cell lysate (WCL) was reserved from a representative replicate and subjected to immunoblotting as described below. To account for potential differences in transfection efficiency, firefly luciferase activity was normalized to *Renilla* luciferase values and plotted as fold activity calculated relative to the no L control. Mean ± standard error of the mean (SEM) values and paired, two-tailed t tests were calculated using GraphPad Prism 7.05 software.

## Results

### EBOV VP35 homo-oligomerization domain contains a putative regulatory serine phosphorylation site

Given that VP35 is analogous to the phosphoprotein of rhabdoviruses and paramyxoviruses, we first sought to examine EBOV VP35 for sites of phosphorylation by mass spectrometry (MS). Affinity-tagged VP35 was over-expressed in HeLa cells and isolated by immunoaffinity purification. Among predicted phosphorylation sites identified by MS analysis, Ser129 was chosen to evaluate for regulatory phosphorylation function due to its position, which lies just outside the coiled-coil region responsible for VP35 homo-oligomerization (Figure 1A and C). Moreover, multiple sequence alignment of all ebolavirus species revealed conservation among all ebolavirus species except BOMV (Figure 1B). BOMV is a newly identified ebolavirus species and has not yet been shown to be pathogenic in humans. All other ebolaviruses except RESTV are pathogenic in humans, thus we reasoned the conservation of Ser129 may be functionally important. Accordingly, it was hypothesized that Ser129 plays an important regulatory role for VP35 as a NNSV P protein.

**Figure 1.**
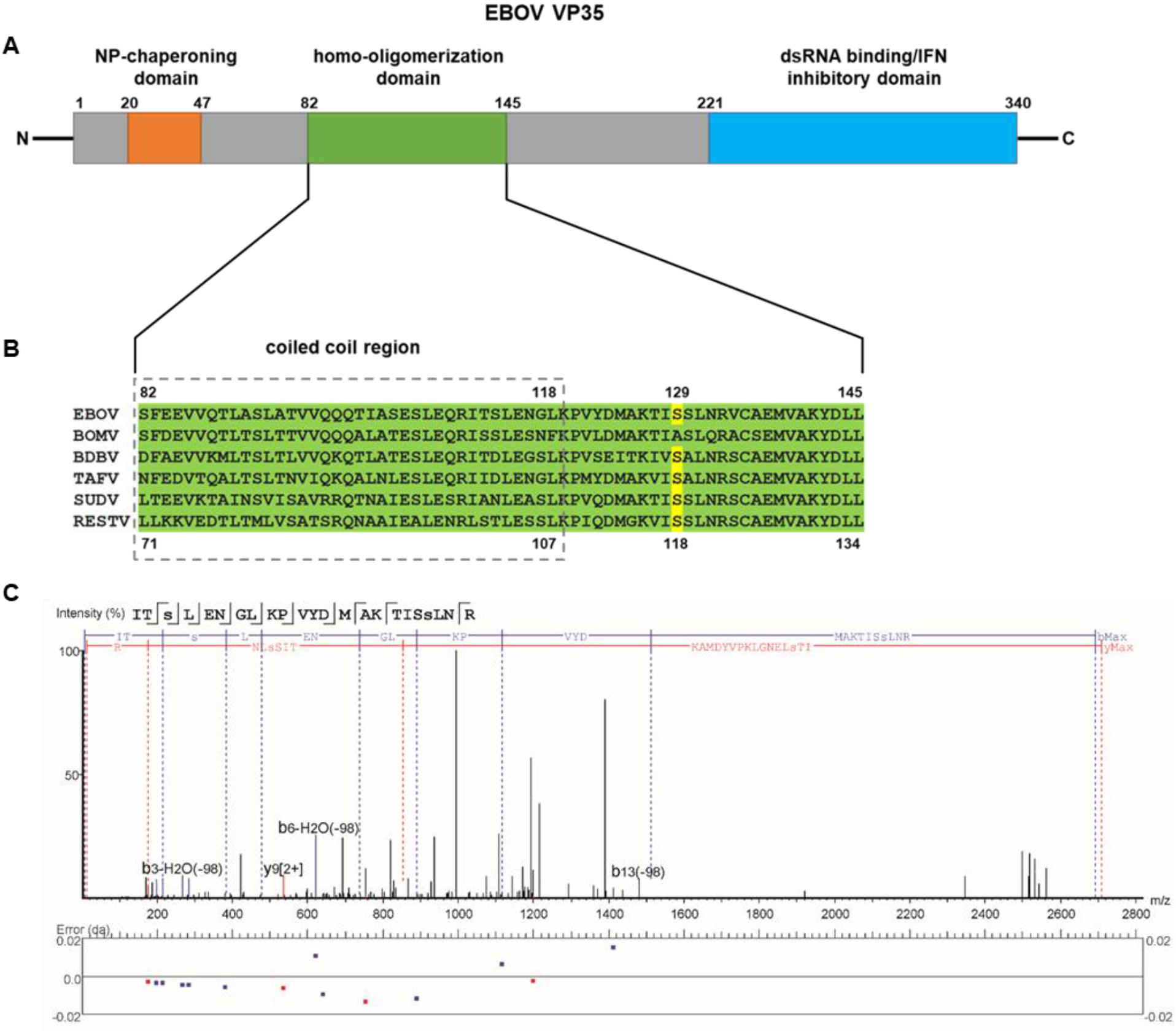
EBOV VP35 homo-oligomerization domain contains a putative regulatory serine phosphorylation site. (A) A schematic representation of VP35 domain organization. (B) Multiple sequence alignment of the homo-oligomerization domain of all ebolavirus species. Dashed gray box indicates the coiled coil region. Ser129 is highlighted in yellow. (C) MS/MS spectrum of the VP35 peptide 111-133 from affinity purified FLAG-VP35 for PTM prediction. FLAG-VP35 was overexpressed in HeLa cells, immunoprecipitated, resolved on 10% SDS-PAGE, stained with colloidal Coomassie Blue, and in-gel digested with trypsin. Eluted peptides were analyzed by a Thermo Orbitrap Fusion Lumos Tribrid mass spectrometer. The spectrum gives positive identification of ITSLENGLKPVYDMAKTISSLNR with a phosphorylation site predicted for Ser129.

### VP35-S129A has moderately diminished oligomerization capacity

Homo-oligomerization of VP35 is known to be required for both replication and contributes to anti-IFN functions (Reid et al., 2005, Moller et al., 2005). Thus, we sought to determine whether Ser129 impacts the ability of VP35 to oligomerize. Using a non-phosphorylated mimic of VP35 generated by alanine substitution at Ser129 (VP35-S129A), we first evaluated the ability of VP35-S129A to interact with VP35-WT. HeLa cells were transfected with HA-VP35-WT or His-VP35-S129A alone and in combination with FLAG-VP35-WT. After 24 h, proteins were extracted and immunoprecipitations (IPs) using anti-FLAG beads were performed. IP samples and whole cell lysates (WCLs) were then subjected to SDS-PAGE and products were analyzed by immunoblot. As expected, HA-VP35-WT was shown to be pulled down only in the presence of FLAG-VP35-WT, demonstrating specificity of binding (Figure 2A, top panels). Similarly, His-VP35-S129A was also pulled down in the presence of FLAG-VP35-WT, indicating that the VP35-S129A mutant retains the ability to self-associate. Appropriate protein expression was confirmed by WCL analysis (Figure 2A, bottom panels).

**Figure 2.**
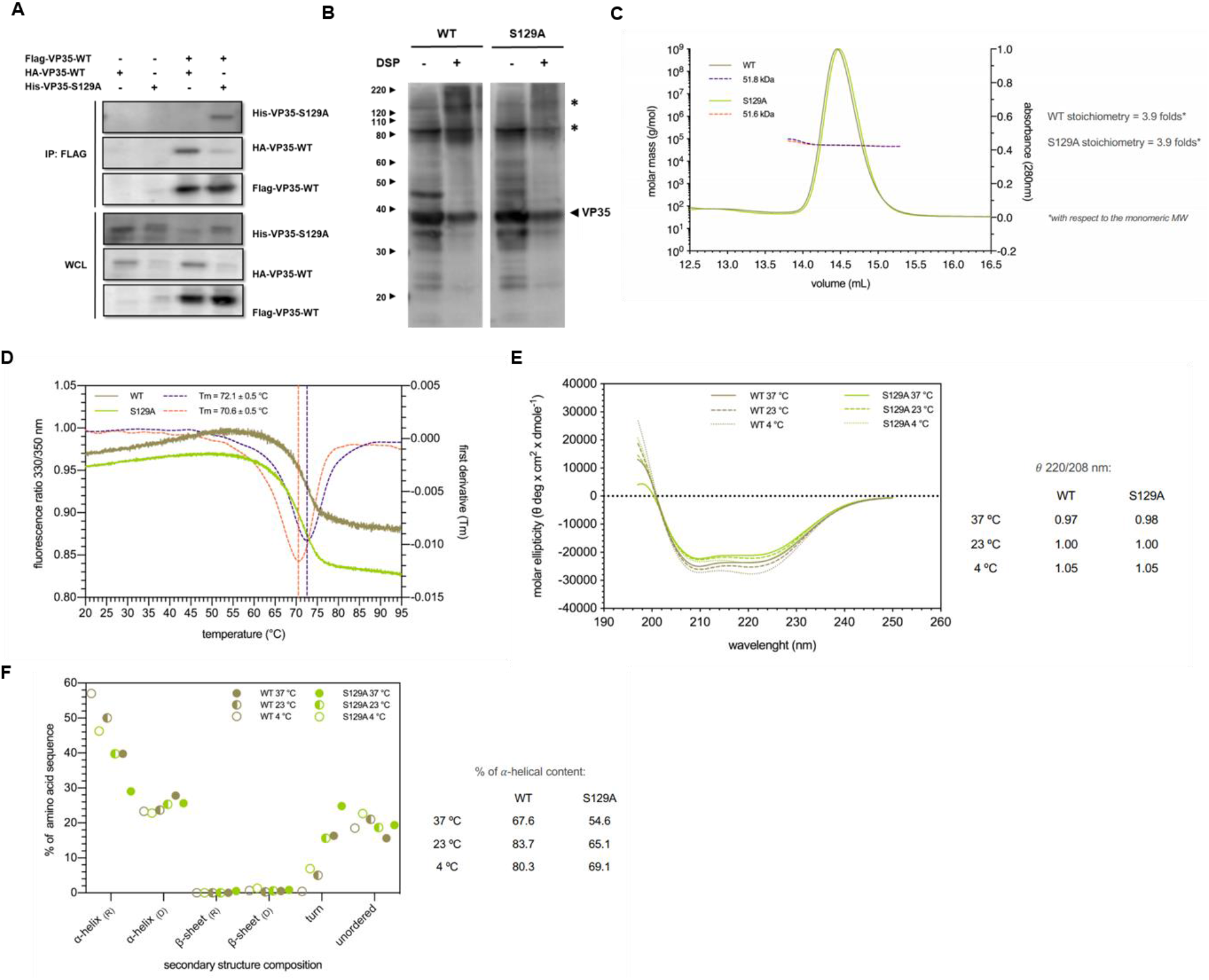
VP35-S129A has moderately diminished oligomerization capacity. (A) CoIP experiment demonstrating that VP35-S129A retains the ability to interact with VP35-WT. IP was performed with anti-FLAG beads after expression of indicated plasmid in HeLa cells. Vector was used to keep amount of DNA transfected constant. (B) DSP cross-linking experiment demonstrating that VP35-S129A retains the capacity to oligomerize at levels near that of WT. HeLa cells were transfected with VP35-WT or VP35-S129A. Twenty-four h after transfection, cells were left untreated or treated with 1 mM of DSP. Asterisks denote oligomeric forms. (C) Quaternary structure analysis of EBOV VP35-WT oligomerization domain compared to S129A mutant by SEC-MALS. Absorbance peaks (280 nm) of protein ertention volumes and absolute molecular masses under the main peak of each sample are indicated by continuous and dashed lines, respectively. (D) Thermal stability analysis of EBOV VP35-WT and S129A oligomerization domain by nanoDSF. Variation in the protein intrinsic 330/350 nm fluorescence ratio upon thermal denaturation and the corresponding first derivative curves are indicated by continuous and dashed lines, respectively. Tm values are defined by the first derivative curve peak of each sample. (E) Far-UV CD spectra of EBOV VP35-WT and S129A oligomerization domain at different temperature values. The 222/208 nm MRE ratio value indicating the coiled-coil folding of each sample is annotated. (F) Secondary structure content analysis of EBOV VP35-WT and S129A oligomerization domain by CD spectra deconvolution with the CONTINLL method. Structure composition of *α*-helices are indicated as percentages of the entire amino acid sequence.

To next evaluate VP35-S129A oligomerization ability, we performed cross-linking experiments, along with a series of biochemical studies. VP35-WT and VP35-S129A were separately over-expressed in HeLa cells for 24 h. Following cross-linking with Dithiobis(succinimidylpropionate) (DSP), the oligomeric forms of VP35 were analyzed by immunoblot under non-reducing, unboiled conditions. After cross-linking with DSP, VP35-WT had a marked increase in higher oligomeric forms, as well as a modest decrease in monomeric form, relative to that of the non-DSP treated VP35-WT isolate (Figure 2B). Notably though, cross-linking of VP35-S129A resulted in a modest reduction in the formation of higher oligomeric forms compared to VP35-WT. This indicates that while decreased, VP35-S129A can still oligomerize.

Full length VP35 has been shown to form both trimers and tetramers in solution (Reid et al., 2005, Zinzula et al., 2009, Luthra et al., 2015, Bruhn et al., 2017, Ramaswamy et al., 2018), with the homo-tetrameric state being the dominant one among ebolavirus species (Zinzula et al., 2019b, Chanthamontri et al., 2019). Given that our VP35-S129A mutant displayed reduced capability to undergo oligomerization under cross-linking experiments, we next wanted to assess if the introduced mutation was affecting the quaternary structure of the oligomerization domain. To this aim, we performed size-exclusion chromatography coupled with multi-angle light scattering (SEC-MALS) on a truncated VP35 construct bordering the N-terminal coiled-coil (residues 75-185). WT and S129A VP35 oligomerization domains both eluted in one distinct peak with an average molecular mass of 51.8 and 51.6 kDa, respectively (Figure 2C). Corresponding to roughly 3.9 folds in stoichiometry compared to their monomeric molecular weight, these masses are consistent with the presence of a homo-tetramer in solution for both WT and S129A VP35. We then wondered if the defective oligomerization phenotype observed by the cross-linking immunoblot was due to any difference in stability between the homo-tetrameric oligomerization domain of the WT and the S129A mutant. To address this question, we performed thermal stability analysis by miniaturized differential scanning fluorimetry (nanoDSF). As shown by their thermal denaturation curves, WT and S129 VP35 oligomerization domains displayed melting temperature (Tm) values of 72.1°C and 70.6°C, respectively (Figure 2D). Although both these inflection points are reminiscent of a stable tertiary structure, the lower Tm displayed by the S129A mutant suggests that a higher degree of flexibility exists at the level of the VP35 oligomerization domain.

Therefore, in order to assess if the S129A mutation could cause structural perturbations sufficient to locally affect the oligomerization domain folding, we tested it by circular dichroism (CD) spectroscopy. In agreement to what has been previously observed for the WT VP35 oligomerization domain (Luthra et al., 2015; Zinzula et al., 2019), the CD spectra of the VP35 S129A mutant was typical to that of a protein mainly consisting of *α*-helices. Moreover, the value of the 222/208 nm ellipticity ratio was close to 1 for both proteins at all three temperature values tested (4°C, 23°C and 37°C), indicating that the coiled-coil motif - through which the *α*-helices interact – remains in place in the S129A mutant. However, the difference in the ellipticity profile of S129A suggests that this mutation introduces some local conformational change (Figure 2E). Consistent with this hypothesis, deconvolution of CD spectra and analysis of the secondary structure content showed that the VP35 S129A oligomerization domain has a lower *α*-helical percentage with respect to WT at all temperature values tested, and that this content diminishes as temperature increases (Figure 2F). This result indicates that a more flexible and relaxed conformation is applied at the level of the VP35 coiled-coil superhelix upon the substitution of Ser-to-Ala at residue 129.

We further investigated the effect of S129A on VP35 oligomerization domain conformation by computational modeling. Starting from the available WT crystal structure, a Ser-to-Ala substitution was made and the oligomerization domain was subjected to energy minimization. The resulting structure was then compared to WT (Figure 3A and B). Conformation was unaffected but did cause a loss of hydrogen bonds between Ser129 and the neighboring Ala125 (Figure 3C). The implicated biological effect may contribute to protein-protein interaction or modulatory phosphorylation (Figure 3D). Taken together, these data indicate that VP35-S129A has reduced capacity to oligomerize relative to VP35-WT and suggests a biological effect.

**Figure 3.**
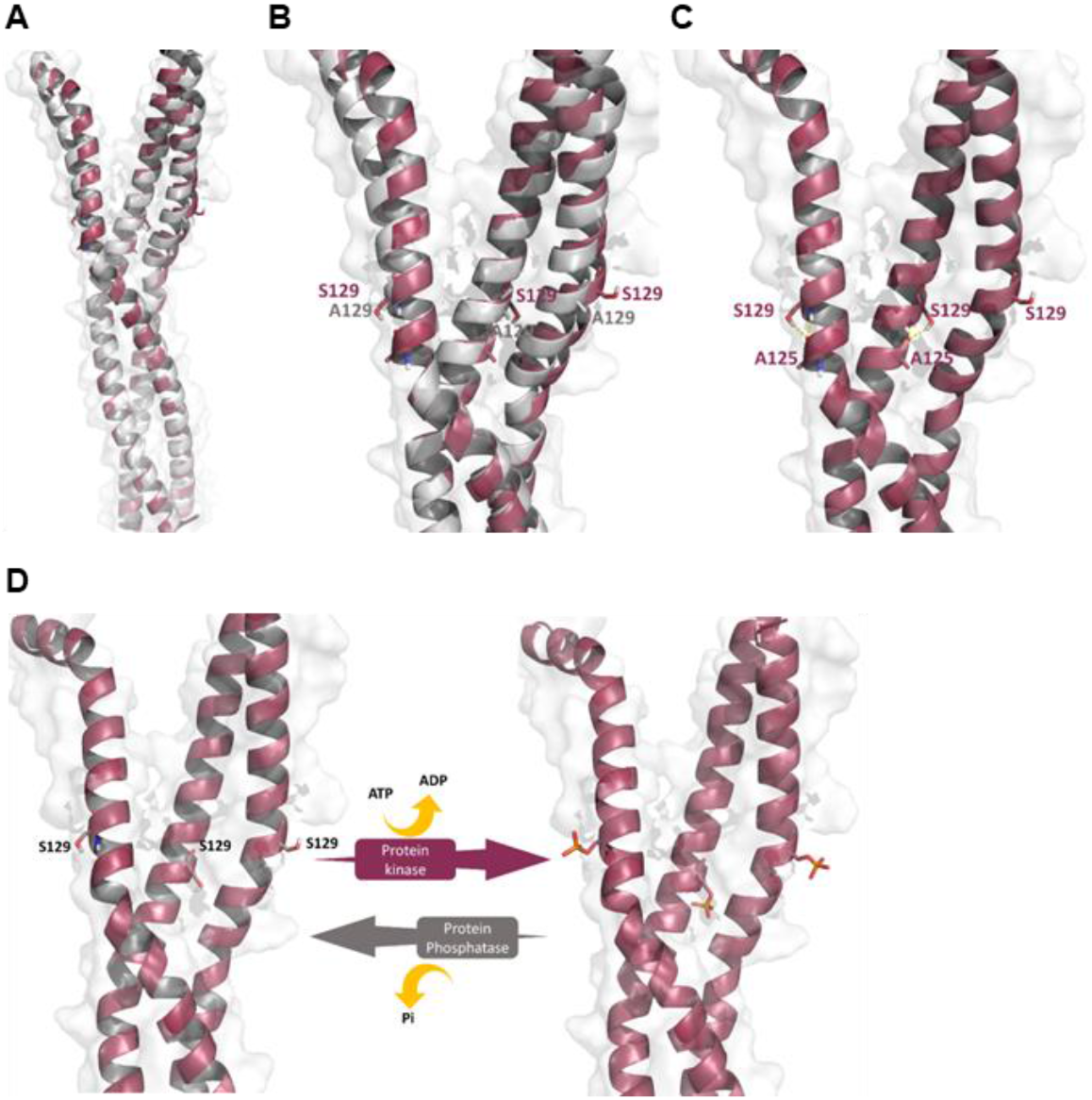
Predicted phosphorylation at Ser129 of VP35 oligomerization domain implicates biological effect. (A) Comparison by computational modeling of VP35-WT (maroon) and VP35-S129A (grey) oligomerization domain. (B) Close-up. (C) Hydrogen bond network. (D) Phosphorylation at Ser129 of VP35 oligomerization domain.

### VP35-S129A retains IFN antagonist function

To assess a role of Ser129 in VP35 function, we examined IFN antagonist function, including VP35 inhibition impact on the RIG-I–mediated signaling cascade and subsequent impairment of the interferon stimulated-gene (ISG) expression. Using a luciferase reporter gene assay, we first evaluated VP35-WT and VP35-S129A inhibition of dsRNA-induced RIG-I activation of the IFN-β promoter. HEK 293T cells were co-transfected with pGL-IFN-β-luc, pRL-TK, and various dilutions of a vector control, VP35-WT, or VP35-S129A. Twenty-four h post-transfection, cells were stimulated with influenza A virus (IAV)-RNA. After 24 h, reporter activity was measured to assess IFN inhibition. IFN-β promoter activation was significantly inhibited by both VP35-WT and VP35-S129A at comparable efficiencies (Figure 4A). Next, we tested the effect of VP35-WT and VP35-S129A on expression of ISGs upon stimulation by viral RNA or poly I:C. Using RT-qPCR, we analyzed gene expression of two well characterized ISGs, ISG15 and 2’-5’-oligoadenylate synthetase (OAS 2’-5’). Expression of these ISGs induced by stimulation with vRNA was significantly reduced by the presence of either VP35-WT or VP35-S129A, relative to the vector control. Specifically, ISG15 expression was inhibited 87% by VP35-WT and 90% by VP35-S129A, and OAS 2’-5’ expression was inhibited 87% by VP35-WT and 95% by VP35-S129A (Figure 4B). Further, the effect of poly I:C stimulation on ISG56 protein expression was assessed by immunoblot. HeLa cells were transfected with vector control, VP35-WT, or VP35-S129A. Twenty-four h post-transfection cells were left unstimulated or stimulated with poly I:C.

**Figure 4.**
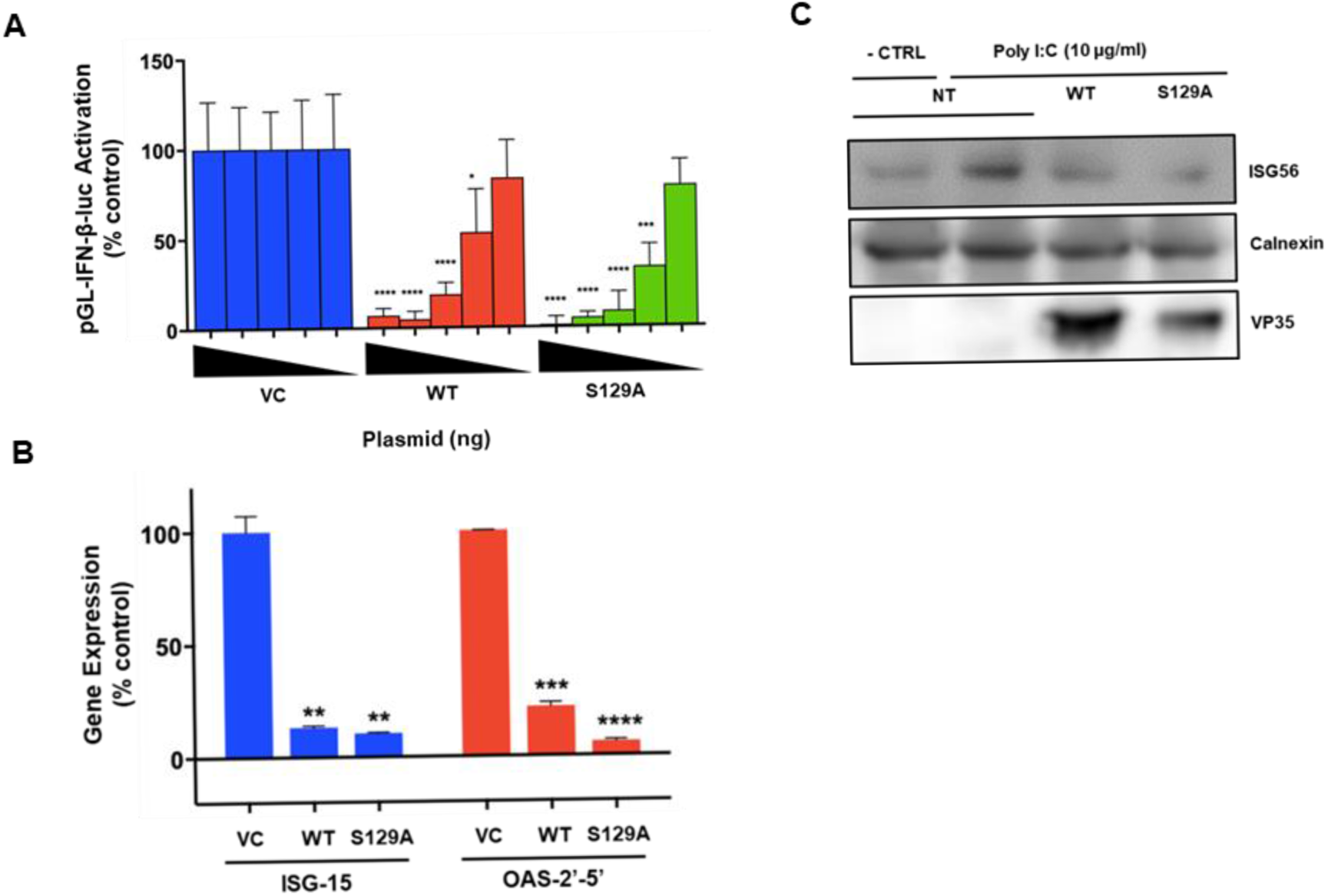
VP35-S129A retains IFN antagonist function. (A) IFN-β promoter activity in the presence of vector control (VC), VP35-WT (WT) or VP35-Ser129Ala (S129A). HEK293T cells were co-transfected with 60 ng of pGL-IFN-β-luc, 10 ng pRL-TK, and decreasing amounts (100, 10, 1.0, 0.1, 0.01 ng) of VC, WT, or S129A plasmid. Twenty-four h after transfection, cells were additionally transfected with IAV-RNA. After 24 h cells were lysed and luciferase activity was measured. (B) RT-qPCR assay of interferon-stimulated gene (ISG) expression upon IAV-RNA stimulation in the presence of VC, WT, or S129A plasmids. HEK293T were transfected with 2500 ng of VC, WT, or S129A plasmid. Twenty-four h after transfection, cells were stimulated with 2500 ng of IAV-RNA. After 24 h, total RNA was extracted, reverse transcribed, and subjected to quantitative real-time PCR (RT-qPCR) for the analysis of ISG15 and 2’-5’-oligoadenylate synthetase (OAS 2’-5’) levels. mRNA expression levels were normalized to the level of glyceraldehyde-3-phosphate dehydrogenase (GAPDH). Data represent mean ± SD (n=3) of two independent experiments. (C) Effect of VP35-WT and VP35-S129A on ISG56 expression upon poly I:C stimulation. HeLa cells were left untransfected or transfected with VP35-WT or VP35-S129A. Twenty-four h after transfection, cells were left untreated or treated with 10 μg/mL of poly I:C. After 24 h, cells were harvested and subjected to immunoblotting. Data represent mean ± SD. *p ≤ 0.05, **p ≤ 0.01, *** p ≤ 0.001, **** ≤ 0.0001.

Both VP35-WT and VP35-S129A were able to inhibit ISG56 protein expression, with levels comparable to that of the non-stimulated vector control (Figure 4C). Collectively, the data show that VP35-S129A retains IFN antagonist activity, indicating that VP35 Ser129 does not impact IFN antagonist function.

### VP35-S129A abrogates EBOV minigenome activity and interaction with L_1-505_

Since VP35 is an essential polymerase cofactor, we next evaluated the effect of VP35-S129A on EBOV minigenome activity. Plasmids encoding EBOV minigenome assay components NP, L, VP35, VP30 and the 3E5E-luciferase minigenome, along with *Renilla* luciferase to control for transfection efficiency variability, were co-transfected in HeLa cells. Forty-eight h post-transfection, a dual-luciferase assay was used to measure reporter activity. Notably, the presence of VP35-S129A nearly abolished minigenome activity relative to VP35-WT, with activity level close to that of the no L control (Figure 5A). Additionally, representative lysates from the minigenome assay were subjected to immunoblotting to confirm appropriate expression of the WT and mutant (Figure 5B). While both VP35-WT and VP35-S129A predominantly expressed one major species of the same migration (~38 kD), other bands were observed. Specifically, VP35-WT contained modest upper bands at ~45 kD and ~85 kD), whereas VP35-S129A contained a modest lower band at ~35 kD. This may be representative of differential PTMs between VP35-WT and VP35-S129A, such as differing phosphorylation states. Overall, these data suggest that Ser129 impacts VP35 transcription and replication function.

**Figure 5.**
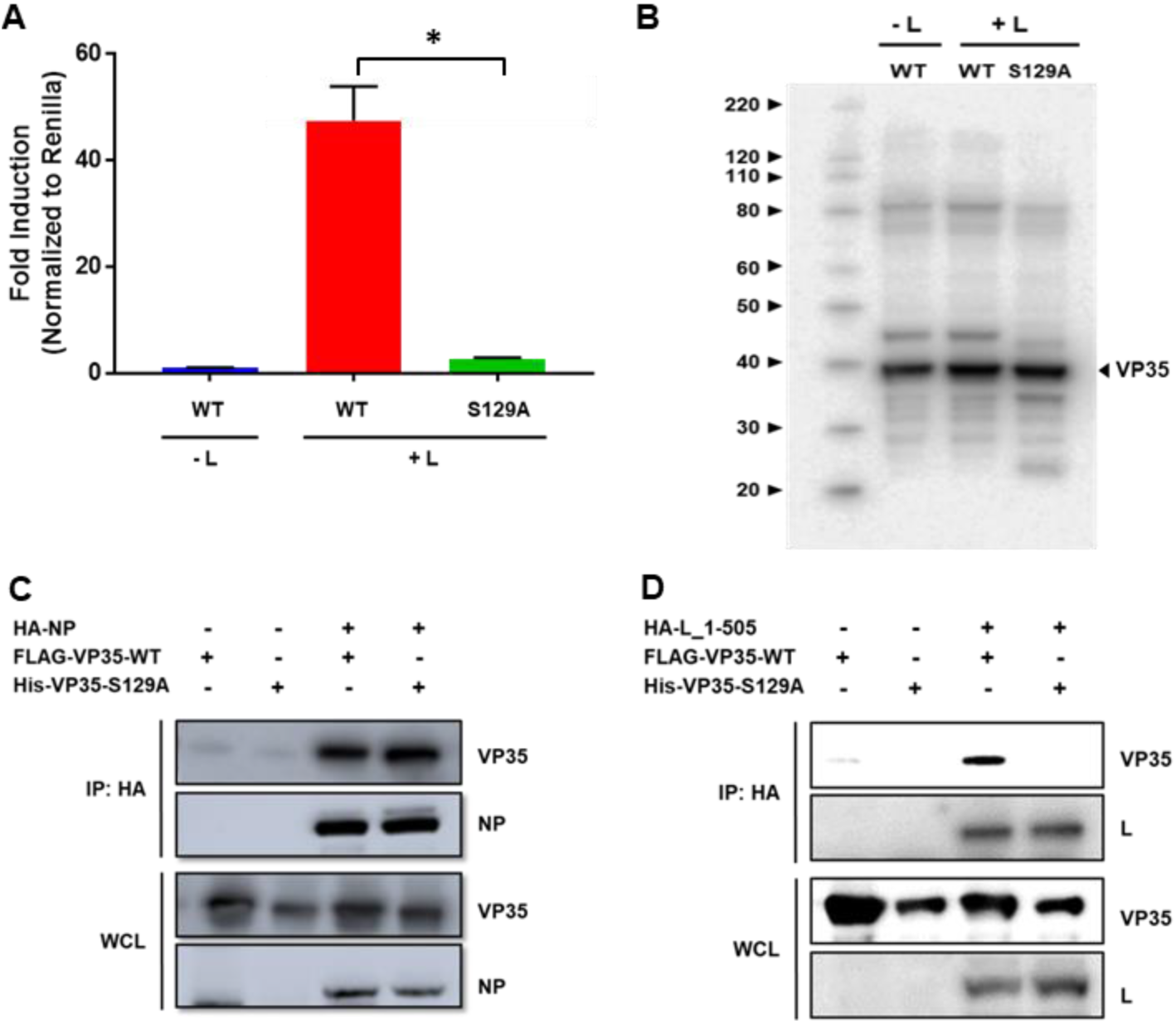
VP35-S129A abrogates EBOV minigenome activity and interaction with L_1-505._ A) Effect of VP35-WT and VP35-S129A on minigenome activity. HeLa cells were transfected with minigenome components (125 ng pCAGGS-HA-NP, 125 ng pCAGGS-FLAG-VP35, 50 ng pCAGGS-V5-VP30, 50 ng pRL-TK, 500 ng pCAGGS-L, and 750 ng of pCAGGS-3E5E-luciferase). Forty-eight h post-transfection, reporter activity was measured. Data represent mean ± SEM from one representative experiment (n=3) of at least three independent experiments (B) Immunoblot confirmation of VP35-WT and VP35-S129A protein levels in the minigenome (C) CoIP experiment demonstrating that VP35-S129A retains the ability to interact with NP. (D) Representative IF images of FLAG-VP35-WT or HIS-VP35-S129A in the presence of HA-NP. (E) CoIP experiment demonstrating a lost interaction between VP35-S129A and L_1-505_. IPs in (C and E) were performed with anti-HA beads. FLAG-VP35-WT or HIS-VP35-S129A (red), HA-NP or HA-L_1-505_, Hoechst 33342 nuclear stain (blue) were visualized by confocal microscopy. Scale bars = 20 uM. *p ≤ 0.05.

To further investigate the role of Ser129 on VP35 replication function, we next examined whether VP35-S129A retains the ability to interact with NP and L by co-IP experiments. HeLa cells were transfected with FLAG-VP35-WT or HIS-VP35-S129A in the absence or presence of either HA-NP or HA-L_1-505_. The previously described HA-L_1-505_ truncation mutant includes the VP35 interaction site and was used due to protein detection issues of full length L (Shabman et al., 2013, Trunschke et al., 2013). Vector control was included in transfections with VP35-WT or VP35-S129A alone to keep total DNA amounts constant. After 24 h, proteins were harvested and applied to IP using HA-tagged beads. After IP, WCLs and IP samples were subjected to SDS-PAGE and products were analyzed by immunoblot. Both VP35-WT and VP35-S129A were pulled down only in the presence of HA-NP, demonstrating specificity of interaction and indicating that VP35-S129A retains the ability to interact with NP (Figure 5C). Notably, VP35-WT was pulled down with HA-L_1-505_ whereas VP35-S129A was not, indicating that the substitution of Ser129 to Ala results in the loss VP35-L interaction (Figure 5D). Collectively, these data show that Ser-to-Ala substitution at residue 129 of VP35 abolishes minigenome activity through loss of interaction between VP35-S129A and L but does not affect IFN antagonist function, effectively uncoupling IFN antagonist and replication functions. This is the first report to suggest phosphorylation contributes a regulatory role in VP35 function and presents a potential therapeutic target.

## Discussion

The most highly phosphorylated protein of NNSVs is generally the P protein, with phosphorylation universally regarded as important to P protein function. While phosphorylation has been shown to modulate the function of several NNSVs including VSV, RSV, CHPV, BDV, RABV, RPV, MV, MuV, and NDV, evidence to support this is lacking for filoviruses (Chattopadhyay and Banerjee, 1987, Barik and Banerjee, 1991, Barik and Banerjee, 1992a, Barik and Banerjee, 1992b, Gao and Lenard, 1995, Gao et al., 1996, Pattnaik et al., 1997, Das and Pattnaik, 2004, Barik et al., 1995, Asenjo and Villanueva, 2000, Villanueva et al., 2000, Asenjo et al., 2006, Asenjo et al., 2008, Asenjo and Villanueva, 2016, Chattopadhyay et al., 1997, Raha et al., 1999, Raha et al., 2000, Schmid et al., 2007, Timani et al., 2008, Moseley et al., 2007, Saikia et al., 2008, Sugai et al., 2012, Pickar et al., 2014, Qiu et al., 2016). The combination of cell-biological, biochemical, and computational studies described here suggests that phosphorylation plays a modulatory role in EBOV VP35 function, thus supporting phosphorylation of the EBOV P protein as functionally significant. Specifically, we identify a highly conserved Ser129 as a key regulatory residue, showing by Ala substitution that Ser129 is important for VP35 replication function, but not IFN antagonist function. NanoDSF results indicate that the S129A mutant possesses a higher degree of flexibility at the level of the VP35 oligomerization domain, with secondary structure content analysis indicating that a more flexible and relaxed conformation is applied at the level of the VP35 coiled-coil superhelix. We further show that while interactions with VP35-WT and NP remain intact, VP35-S129A exhibits impaired ability to interact with the viral polymerase.

VP35, along with L, NP, and VP30, is an essential component of the filovirus replication machinery, functioning as a non-enzymatic cofactor of the viral polymerase (Muhlberger et al., 1998, Muhlberger et al., 1999, Boehmann et al., 2005, Prins et al., 2010). It is generally accepted that VP35 acts as a bridge between L and NP, thereby serving to mediate L interaction with NP-encapsidated template RNAs. Both VP35-L and VP35-NP interactions are thus likely to be essential for viral RNA synthesis (Becker et al., 1998, DiCarlo et al., 2007, Trunschke et al., 2013, Kirchdoerfer et al., 2015, Leung et al., 2015). It has been shown that VSV P protein must undergo phosphorylation-dependent homo-oligomerization to become transcriptionally active (Gao and Lenard, 1995). Consistent with these findings are the nanoDSF results in this study which indicate that the VP35-S129A mutant possesses a higher degree of flexibility at the level of the VP35 oligomerization domain, and severely decreases EBOV minigenome activity. In addition, the oligomerization domain of MuV and hPIV3 P proteins has been shown to have an enhancing effect on the P-L interaction, which further supports the findings here (Choudhary et al., 2002, Pickar et al., 2015).

VP35 has been shown to be phosphorylated by IKKɛ and TBK-1 in vitro, which suggests the possibility that its function may be modulated by these kinases (Prins et al., 2009). In this case VP35 exerts IFN-antagonist function by preventing TBK-1 and/or IKKɛ from activating IRF3/IRF7. It has yet to be determined whether VP35 becomes phosphorylated in EBOV-infected cells, though, and whether that modulates its function. The extent to which VP35 is phosphorylated by these kinases in EBOV-infected cells would be influenced by the extent to which VP35 acts as a decoy substrate for IKKɛ and TBK-1 relative to the extent VP35 merely prevents kinase activation via steric inhibition (Prins et al., 2009). Even so, in our study here the VP35-S129A mutant impaired replication function but did not affect VP35 IFN antagonist function. Given the high virulence of EBOV infection and the multifunctional nature of VP35, it seems unlikely that VP35 phosphorylation by IKKɛ and TBK-1 would detrimentally affect virus replication. Considering then that the IFN antagonism of the VP35-S129A mutant remained intact instead suggests that another host kinase(s) would be involved in the potential phosphorylation of Ser129.

Previous studies have demonstrated phosphorylation of NP and VP30; moreover, a functional significance for VP30 phosphorylation has been described (Elliott et al., 1985, Becker et al., 1994, Modrof et al., 2002, Martinez et al., 2008, Biedenkopf et al., 2016). The data here suggest that phosphorylation of VP35 does play a modulatory role, which aligns with the homologous functions of NNSV P proteins. Even so, the MS analysis herein was limited to predicting phosphorylation of Ser129, rather than definitively identifying the residue as phosphorylated, thus future studies will need to provide such evidence. Given that other P proteins, such as VSV and RSV, have been dramatically affected by single Ser-to-Ala substitutions lends more credence to phosphorylation of VP35 Ser129 exerting a modulatory role, as opposed to other PTMs (Chattopadhyay and Banerjee, 1987, Asenjo and Villanueva, 2000). However, other modifications of Ser are known to occur, including O-linked glycosylation and acetylation (Wang et al., 2015). Future studies must confirm relevance during EBOV infection as well as determine whether Ser129 is post-translationally modified, and, if so, whether the PTM is constitutive or dynamic.

EBOV, SUDV, and BDBV infections cause severe disease in humans with high case fatality rates (Feldmann et al., 2013). While there are promising vaccine and therapeutic candidates, there remains an urgent need to develop effective therapeutics against ebolaviruses (Haque et al., 2015, Espeland et al., 2018, Dhama et al., 2018). Development of drugs that interfere with VP35 homo-oligomerization are likely to impair viral gene expression, therefore identification of a potential target is an asset for the design of novel antiviral agents. Because Ser129 is well conserved across ebolaviruses, inhibition of its modulatory role presents a potential pan-filoviral therapeutic strategy.

In recent years, novel VP35 activities beyond IFN antagonist, polymerase cofactor, and nucleocapsid component have been discovered, including repression of stress granules and NTPase and helicase-like activities (Le Sage et al., 2017, Shu et al., 2019). Though these studies have provided more insight into VP35 biology, the regulation of these activities is underexplored, particularly regarding the homology of VP35 as a P protein. Here, our data indicate that the Ser129 residue is important for VP35 polymerase cofactor function but not for IFN antagonist function, which effectively uncouples the major function of VP35. Biochemical characterization indicated that VP35-S129A has reduced capacity to oligomerize, and coIPs showed that the interaction between VP35 and L1-505 was abolished upon Ala substitution at residue 129. Future studies will address host kinases and incorporate viral infections to shed light on how VP35 function is modulated.

## Acknowledgments

We thank Janice A. Taylor and James R. Talaska of the Advanced Microscopy Core Facility at the University of Nebraska Medical Center for providing assistance with confocal microscopy. The University of Nebraska Medical Center Advanced Microscopy Core Facility receives partial support from the National Institute for General Medical Science (NIGMS) INBRE - P20 GM103427 and COBRE - P30 GM106397 grants, as well as support from the National Cancer Institute (NCI) for The Fred & Pamela Buffett Cancer Center Support Grant-P30 CA036727, and the Nebraska Research Initiative. This publication’s contents and interpretations are the sole responsibility of the authors.

